# Slingshot: Cell lineage and pseudotime inference for single-cell transcriptomics

**DOI:** 10.1101/128843

**Authors:** Kelly Street, Davide Risso, Russell B. Fletcher, Diya Das, John Ngai, Nir Yosef, Elizabeth Purdom, Sandrine Dudoit

## Abstract

Single-cell transcriptomics allows researchers to investigate complex communities of heterogeneous cells. These methods can be applied to stem cells and their descendants in order to chart the progression from multipotent progenitors to fully differentiated cells. While a number of statistical and computational methods have been proposed for analyzing cell lineages, the problem of accurately characterizing multiple branching lineages remains difficult to solve. Here, we introduce a novel method, Slingshot, for inferring multiple developmental lineages from single-cell gene expression data. Slingshot is a uniquely robust and flexible tool for inferring developmental lineages and ordering cells to reflect continuous, branching processes.

## 1 Background

Traditional transcription assays, such as bulk microarrays and RNA sequencing (RNA-Seq), offer a bird’s-eye view of transcription. However, as they rely on RNA from a large number of cells as starting material, they are not ideal for examining heterogeneous populations of cells. Newly-developed single-cell assays can give us a much more detailed picture (Kolodziejczyk et al., 2015). This higher resolution allows researchers to distinguish between closely-related populations of cells, potentially revealing functionally distinct groups with complex relationships (Wagner et al., 2016).

For many systems, there are not clear distinctions between cellular states, but instead there is a smooth transition where individual cells represent points along a continuum or “lineage”. Cells in these systems change states by undergoing gradual transcriptional changes, with progress being driven by an underlying temporal variable. For example, Trapnell et al. (2014) examined the differentiation pattern of skeletal myoblasts, showing that their development into myocytes and mature myotubes follows a continuous lineage, rather than discrete steps. Inference of lineage structure has been referred to as “pseudotemporal reconstruction” and it can help us understand how cells change state and how cell fate decisions are made (Bendall et al., 2014; Campbell et al., 2015; Trapnell et al., 2014). Furthermore, many systems contain lineages that share a common initial state but branch and terminate at different states. These complicated lineage structures require additional analysis to distinguish between cells that fall along different lineages (Ji and Ji, 2016; Setty et al., 2016; Shin et al., 2015).

Several methods have been proposed for the task of pseudotemporal reconstruction, each with their own set of strengths and assumptions. We describe a few popular approaches here; for a more complete review see Bacher and Kendziorski (2016). One of the most well-known methods is Monocle (Trapnell et al., 2014), which constructs a minimum spanning tree (MST) on cells in a reduced-dimensionality space created by independent component analysis (ICA) and orders cells via a PQ tree along the longest path through this tree. The direction of this path and the number of branching events are left to the user, who may examine a known set of marker genes or use time of sample collection as indications of initial and terminal cell states. The methods Waterfall (Shin et al., 2015) and TSCAN (Ji and Ji, 2016) instead determine the lineage structure by first clustering cells and then drawing an MST on the cluster centers. Lineages are represented in the low-dimensional space as piecewise linear paths through the tree, providing a simple, unsupervised method for identifying branching events. The output pseudotime values are calculated by orthogonal projection onto these paths, with the identification of the direction and of the cluster of origin again left to the user. The method of Wishbone (Setty et al., 2016), an extension of Wanderlust (Bendall et al., 2014), uses an ensemble of *k*-nearest neighbors (kNN) graphs on cells and a randomly selected group of waypoints to robustly determine cell-to-cell distances. Given a user-specified initial cell, these distances are used to determine the presence or absence of a bifurcation event, which limits the method to only two lineages. Finally, other approaches use smooth curves to represent lineages. For example, Embeddr (Campbell et al., 2015) uses the principal curves method of Hastie and Stuetzle (1989) to infer lineages in a low-dimensional space obtained by a Laplacian eigenmap. Again, the direction of the curve must be specified by the user and the method is limited to a single lineage. See Table 1 for a summary of existing methods.

**Table 1:**
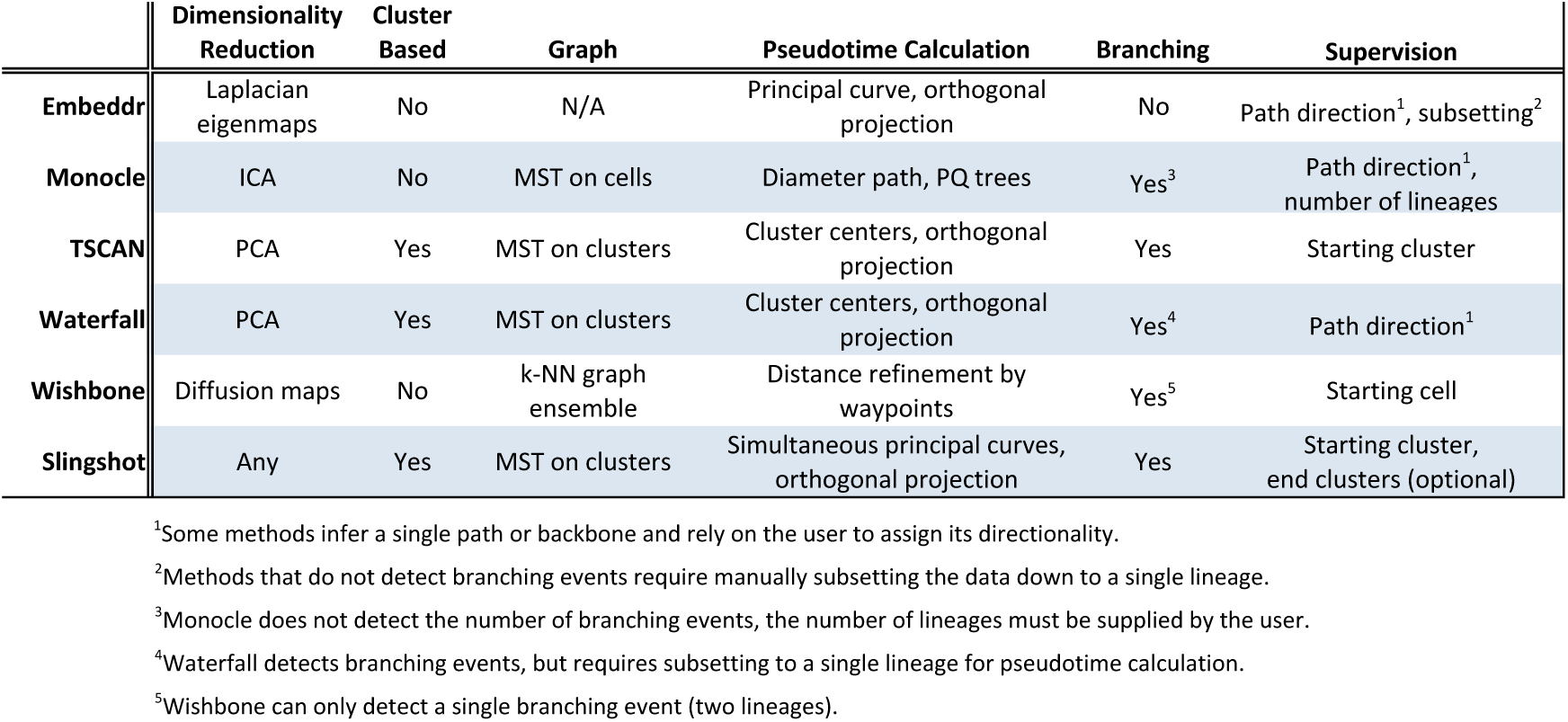
Summaries of existing lineage and pseudotime inference methods within a common framework.

|  | Dimensionality Reduction | Cluster Based | Graph | Pseudotime Calculation | Branching | Supervision |
| --- | --- | --- | --- | --- | --- | --- |
| **Embeddr** | Laplacian eigenmaps | No | N/A | Principal curve, orthogonal projection | No | Path direction<sup>1</sup>, subsetting<sup>2</sup> |
| **Monocle** | ICA | No | MST on cells | Diameter path, PQ trees | Yes<sup>3</sup> | Path direction<sup>1</sup>, number of lineages |
| **TSCAN** | PCA | Yes | MST on clusters | Cluster centers, orthogonal projection | Yes | Starting cluster |
| **Waterfall** | PCA | Yes | MST on clusters | Cluster centers, orthogonal projection | Yes<sup>4</sup> | Path direction<sup>1</sup> |
| **Wishbone** | Diffusion maps | No | k-NN graph ensemble | Distance refinement by waypoints | Yes<sup>5</sup> | Starting cell |
| **Slingshot** | Any | Yes | MST on clusters | Simultaneous principal curves, orthogonal projection | Yes | Starting cluster, end clusters (optional) |

<sup>1</sup>Some methods infer a single path or backbone and rely on the user to assign its directionality.
<sup>2</sup>Methods that do not detect branching events require manually subsetting the data down to a single lineage.
<sup>3</sup>Monocle does not detect the number of branching events, the number of lineages must be supplied by the user.
<sup>4</sup>Waterfall detects branching events, but requires subsetting to a single lineage for pseudotime calculation.
<sup>5</sup>Wishbone can only detect a single branching event (two lineages).

Here, we introduce Slingshot, a novel lineage inference tool designed for multiple, branching lineages. Slingshot combines highly stable techniques necessary for noisy single-cell data with the flexibility to identify multiple lineages with varying levels of supervision. Slingshot consists of two main stages: 1) the inference of the lineage structure and 2) the inference of pseudotime variables for cells along each lineage (Figure 1). Like other methods (Ji and Ji, 2016; Shin et al., 2015), Slingshot’s first stage uses a cluster-based MST to stably identify the key elements of the global lineage structure, i.e., the number of lineages and where they branch (Figure 1, Step 1). This allows us to identify novel lineages while also accommodating the use of domain-specific knowledge to supervise parts of the tree. For the second stage, we propose a novel method, simultaneous principal curves, to fit smooth, branching curves to these lineages, thereby translating the knowledge of global lineage structure into stable estimates of the underlying cell-level pseudotime variable for each lineage (Figure 1, Step 2). The Slingshot method is implemented in the open-source R package slingshot (available from the GitHub repository https://github.com/kstreet13/slingshot) to be released through the Bioconductor Project (http://www.bioconductor.org).

**Figure 1:**
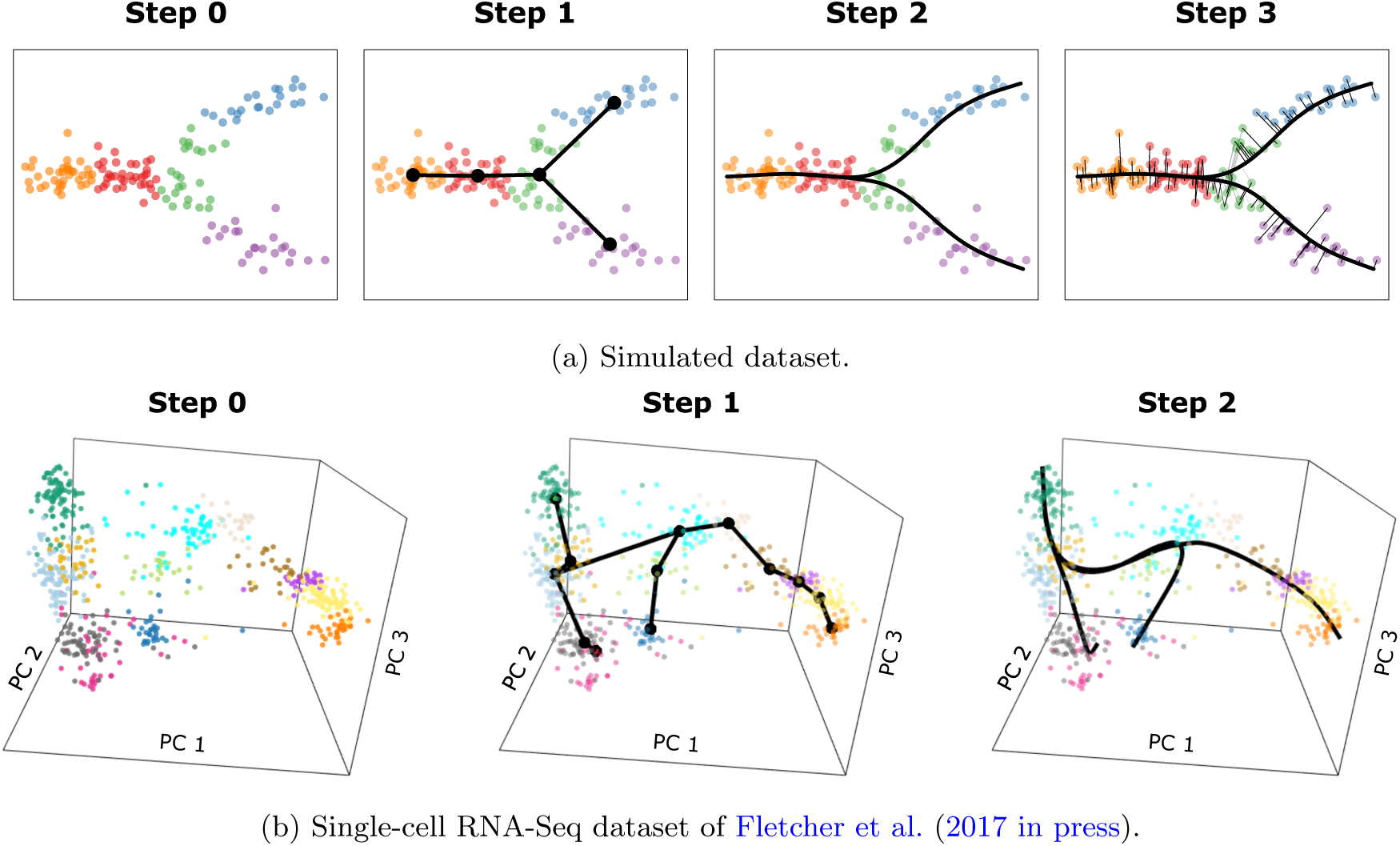
Schematics of Slingshot’s main steps. The main steps for Slingshot are shown for: **Panel** a simple simulated dataset in two dimensions and **Panel (b)** the dataset of Fletcher et al. (2017 in press), discussed in Section 2. Step 0: Slingshot starts from clustered data in a low-dimensional space (cluster labels indicated by color). For **Panel (b)**, the plot shows the top three principal components, but Slingshot was run on the top five. Step 1: A minimum spanning tree is constructed on the clusters to determine the number and rough shape of lineages. For **Panel (b)**, we impose some constraints on the MST based on known biology, see Section 2. Step 2: Simultaneous principal curves are used to obtain smooth representations of each lineage. Step 3: Pseudotime values are obtained by orthogonal projection onto the curves (only shown for **Panel (a)**).

In addition to Slingshot’s core methodological components described above for lineage and pseudotime inference, we note the importance of upstream analysis choices. Indeed, many pseudotemporal inference methods will either implicitly or explicitly require certain choices at previous steps in the workflow. Dimensionality reduction, for example, helps in reducing the amount of noise in the data and in visualizing the data, but a variety of approaches are available, with a potentially large impact on the final result (see Figure S1). Monocle uses independent component analysis, Waterfall and TSCAN use principal component analysis (PCA), Embedder uses Laplacian eigenmaps (Belkin and Niyogi, 2003), and Wishbone uses diffusion maps for analysis and t-distributed stochastic neighbor embedding (t-SNE) (Maaten and Hinton, 2008) for visualization (Table 1). Given the great diversity of data being generated by single-cell assays, it seems unlikely that there is a one-size-fits-all solution to the dimensionality reduction problem. Similar issues arise for normalization and clustering methods. These data analysis steps are very important and because different methods hard-code different choices, the methods can be difficult to compare. Slingshot does not specify these crucial upstream choices, but is instead designed with flexibility and modularity in mind, to easily integrate with the normalization, dimensionality reduction, and clustering methods deemed most appropriate for any particular dataset.

## 2 Results

Slingshot divides the problem of multiple lineage inference into two stages: **1.** Identification of *lineages*, i.e., ordered sets of cell clusters, where all lineages share a starting cluster and each leads to a unique terminal cluster. This is achieved by constructing an MST on clusters of cells. **2.** For each lineage, identification of *pseudotime*, i.e., a one-dimensional variable representing each cell’s transcriptional progression toward the terminal state. This is achieved by a method which extends principal curves (Hastie and Stuetzle, 1989) to the case of multiple branching lineages. The primary benefits of Slingshot are robustness and the ability to detect complicated multiple-lineage structures.

### 2.1 Robustness

One of the main difficulties of single-cell RNA-Seq is the high level of noise. In addition to the host of biological and technical confounders that can affect any (bulk) RNA-Seq experiment, single-cell data may contain effects from transcriptional bursting (Chubb et al., 2006; Raj et al., 2006) and drop-out (Kharchenko et al., 2014). Furthermore, downstream analyses such as lineage inference may be affected by upstream computational choices such as normalization and clustering methods. For these reasons, we believe that robustness to noise, unwanted technical effects, and choice of pre-processing methods should be important characteristics of a lineage inference method.

#### Robustness to Noise

We first examined the stability of a few well-known methods using a subset of the HSMM dataset of Trapnell et al. (2014) chosen to represent a single lineage. In Figure 2, we illustrate each method’s ordering of the full set of 212 cells and show how consistently it orders cells over 50 bootstrap subsamples. The Monocle procedure, which constructs an MST on individual cells and orders them according to PQ trees along the diameter path of the MST, was the least stable of the methods we compared. The diameter path drawn by Monocle was highly variable and sensitive to even small amounts of noise; this instability has been previously discussed in Ji and Ji (2016). In contrast, other methods which emphasize stability in the construction of their primary trajectory and obtain pseudotime values based on orthogonal projection, produced much more stable orderings.

**Figure 2:**
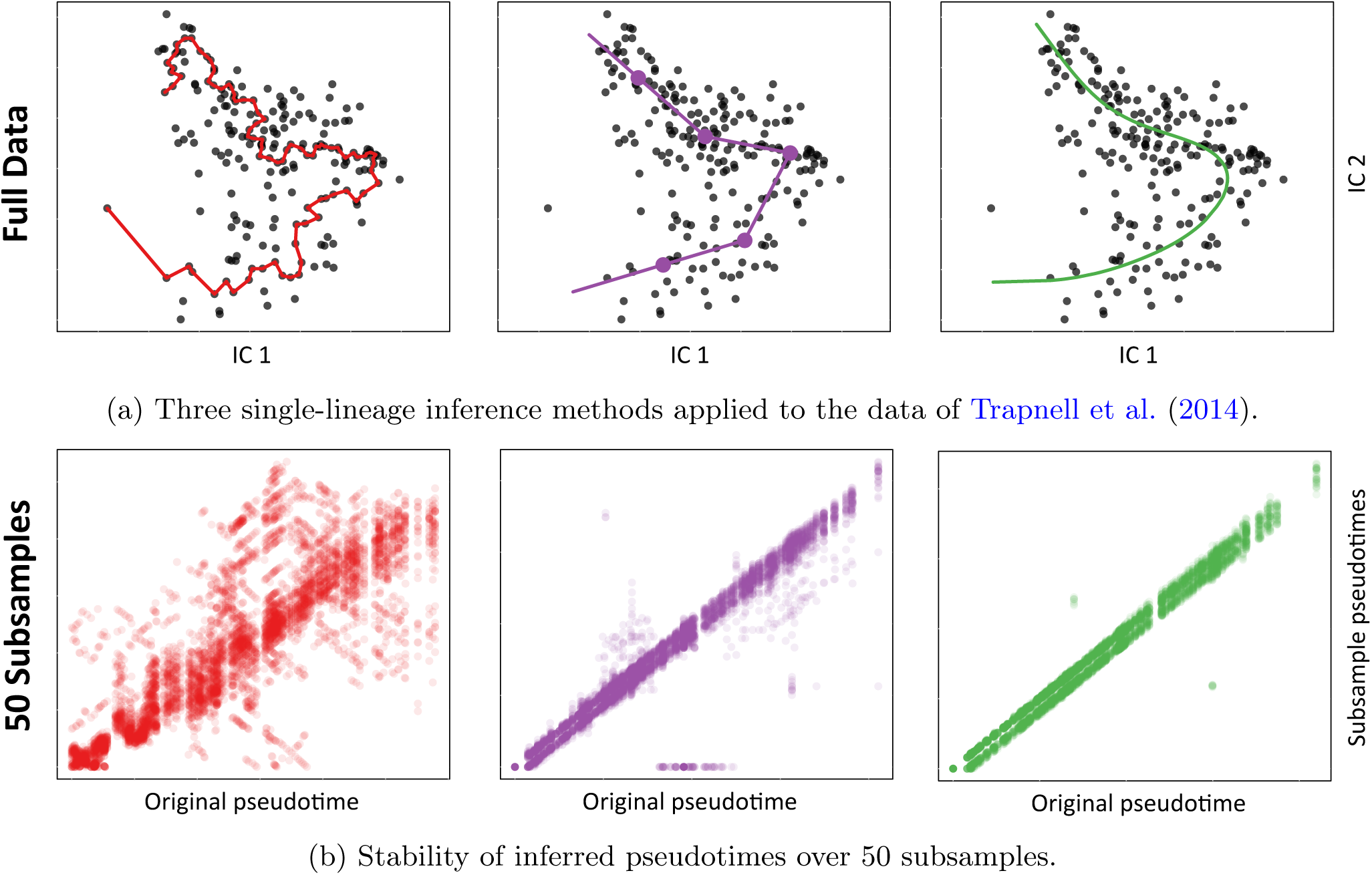
Pseudotime inference: Robustness to noise. We examine the stability of three lineage inference methods, showing how each method orders the cells for the original dataset as well as for 50 subsamples of the data. **Panel (a):** Monocle identifies the longest path through the MST constructed on all cells (red). Waterfall and TSCAN cluster cells and connect cluster centers with an MST (purple, clustering performed by *k*-means with *k* = 5). A principal curve is a non-linear fit through the data, used by Embeddr and Slingshot (green). As in Trapnell et al. (2014), dimensionality reduction is performed by ICA. **Panel (b):** We examine the stability of these three methods by plotting pseudotimes based on 50 subsamples of the data vs. the original pseudotimes. Subsamples were generated in a bootstrap-like manner, by randomly sampling *n* times with replacement from the original cell-level data and retaining only one instance of each cell. Thus, subsamples were of variable sizes, but contained on average about 63% of the original data. The cluster-based MST method occasionally detected spurious branching events and, for the purposes of visualization, cells not placed along the main lineage were assigned a pseudotime value of 0.

Although both the cluster-based MST method (Ji and Ji, 2016; Shin et al., 2015) and the principal curves method (Campbell et al., 2015; Hastie and Stuetzle, 1989) demonstrated robustness over the bootstrap subsamples shown in Figure 2b, the former has some important drawbacks. Due to the sharp corners of the piecewise linear paths, multiple cells will often be assigned identical pseudotime values, corresponding to the value at the vertex. Additionally, the tree does not always capture the correct global structure: due to variability in the exact distances between cluster centers, the MST may end up cutting corners in a lineage and curving back on itself or detecting spurious branching events (see Supplemental Figure S2). The principal curve approach was the most stable method, producing very similar orderings on all subsets of the data (Figure 2b). On more complex datasets, however, Embeddr’s (Campbell et al., 2015) principal curves method has the obvious limitation of only characterizing a single lineage. It is for this reason that we chose to extend principal curves to accommodate multiple, branching lineages.

#### Robustness to Cluster Assignments

Although Slingshot uses cluster assignments for the identification of lineages and branching events, its use of simultaneous principal curves to estimate the final pseudotimes makes its final results quite robust to the choice of clustering method. In Figure 3, we demonstrate on a simulated dataset the stability of the pseudotime estimates for different sets of clusters, chosen by *k*-means clustering. Slingshot produces very similar curves over a wide range of values for the number of clusters *k*, even while the underlying MSTs change dramatically (Figure 3a). In comparison, using an MST alone to identify lineages and estimate pseudotimes (as is done by Waterfall and TSCAN) gives estimates that are quite sensitive to the choice of clusters.

**Figure 3:**
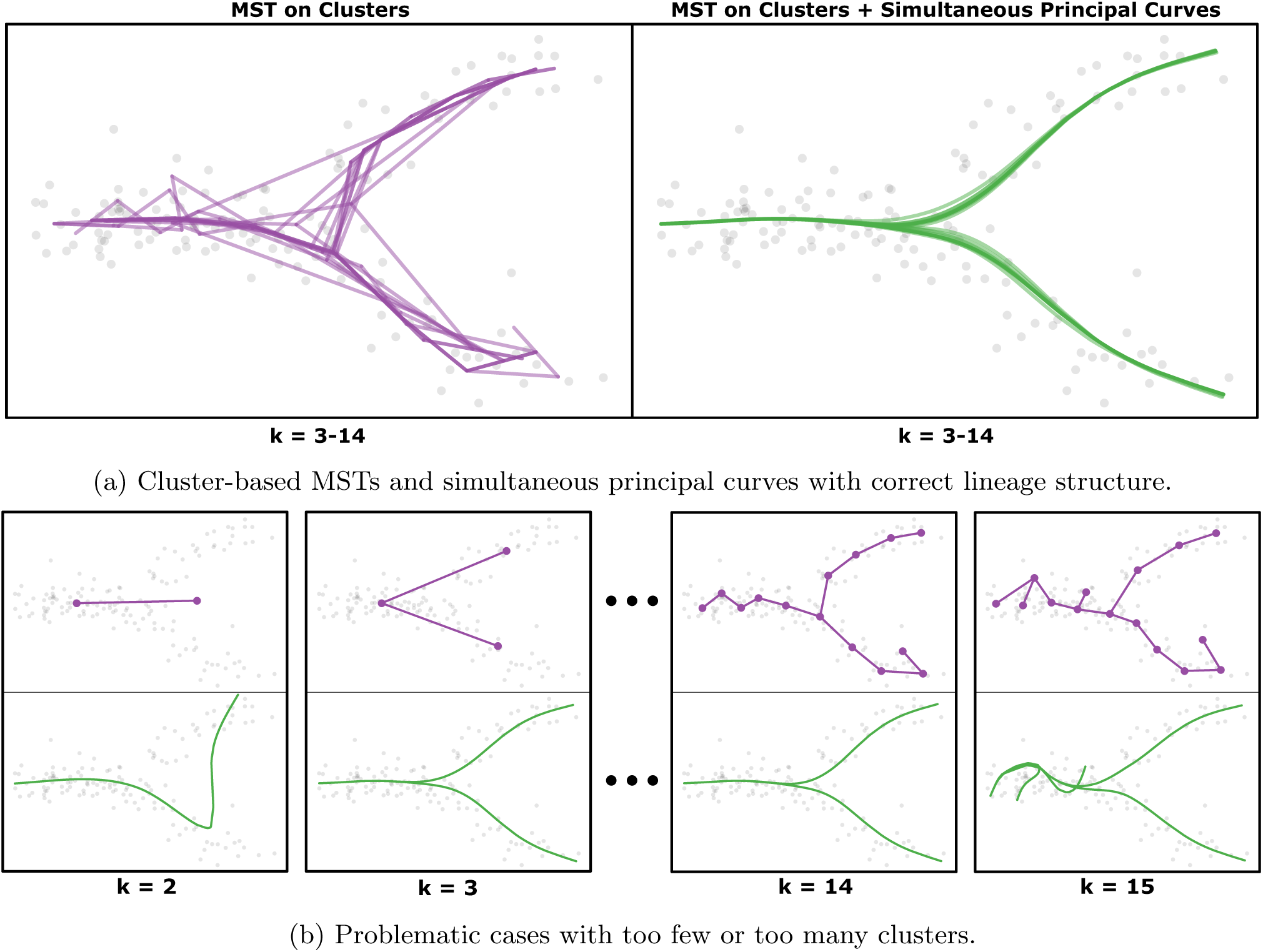
Pseudotime inference: Robustness to cluster assignments. **Panel (a):** Using a simulated two-dimensional dataset with two endpoints, *k*-means to assign cluster labels (*k* = 3*, …,* 14), and Slingshot’s covariance-scaled distance measure, we show that the cluster-based MST can be highly variable, even while identifying the correct global structure (purple, as in TSCAN and Waterfall). Despite using these same trees, Slingshot’s simultaneous principal curves are robust to this variability and all of them produce nearly identical results (green). **Panel (b):** With the same dataset as above, we show two pathological scenarios in which the cluster-based MST fails to identify the correct global structure; the corresponding simultaneous principal curves are plotted in green. Setting *k* = 2, we are unable to detect a branching event and the resulting curve attempts to fit all of the data. With higher numbers of clusters, we eventually run into the problem of overfitting and the MST detects spurious branching events. Thus, for *k* = 15, the corresponding simultaneous principal curves similarly overfit certain regions of the data.

Of course, Slingshot is not completely independent of the effect of poor clustering since the clusters define the lineages. This is seen in the extreme cases where *k* = 2 and *k* = 15 (Figure 3b), for which the inferred lineages are dramatically wrong because of the choice of *k*. Nonetheless, Slingshot shows a reduced reliance on clustering results and increased stability, leading to more robust pseudotime estimates.

### 2.2 Multiple Lineage Identification

Determining the number and location of branching events is one of the most difficult components of lineage inference. Already faced with potentially noisy, high-dimensional data, we now have to consider the problem of model selection in an extremely large model space. Some methods introduce restrictions on lineage discovery; for example, Monocle requires the user to pre-specify the number of lineages and Wishbone is limited to only one or two lineages. In contrast, Slingshot allows for multiple lineage detection without limiting or pre-specifying the number of lineages.

Moreover, Slingshot provides the user the ability to direct lineage detection in a biologically relevant manner, specifically by giving the option to identify known endpoints of lineages. Using the complex example of lineage detection in the olfactory epithelium data of Fletcher et al. (2017 in press), we demonstrate the importance of this type of supervision, as opposed to supervision which requires specific knowledge of the number of lineages.

Figure 4 displays the result of running Slingshot with and without supervision on the data of Fletcher et al. (2017 in press). For this dataset, clusters corresponding to the neuronal, sustentacular, and microvillous cell types were identified based on known marker genes (Fletcher et al., 2017 in press). These cell types are known to be non-differentiating and should thus be the terminal states of their respective lineages. However, without supervision (Figure 4a), the sustentacular cluster was not identified as an endpoint, yielding results inconsistent with prior knowledge. Selecting the sustentacular cluster as an endpoint led to the recovery of the three lineages reported and validated in the original paper (Figure 4b). We also note that this structure could not have been recovered using standard Euclidean distances between cluster centers (Figure 4c), as in Waterfall and TSCAN. By failing to utilize the shapes of the clusters, the standard Euclidean distance between cluster centers identified a spurious branching event very early on in HBC differentiation. By default, Slingshot uses a shape-sensitive distance measure inspired by the Mahalanobis distance (Mahalanobis, 1936), which scales the distance between cluster centers based on the shared covariance structure of the two clusters.

**Figure 4:**
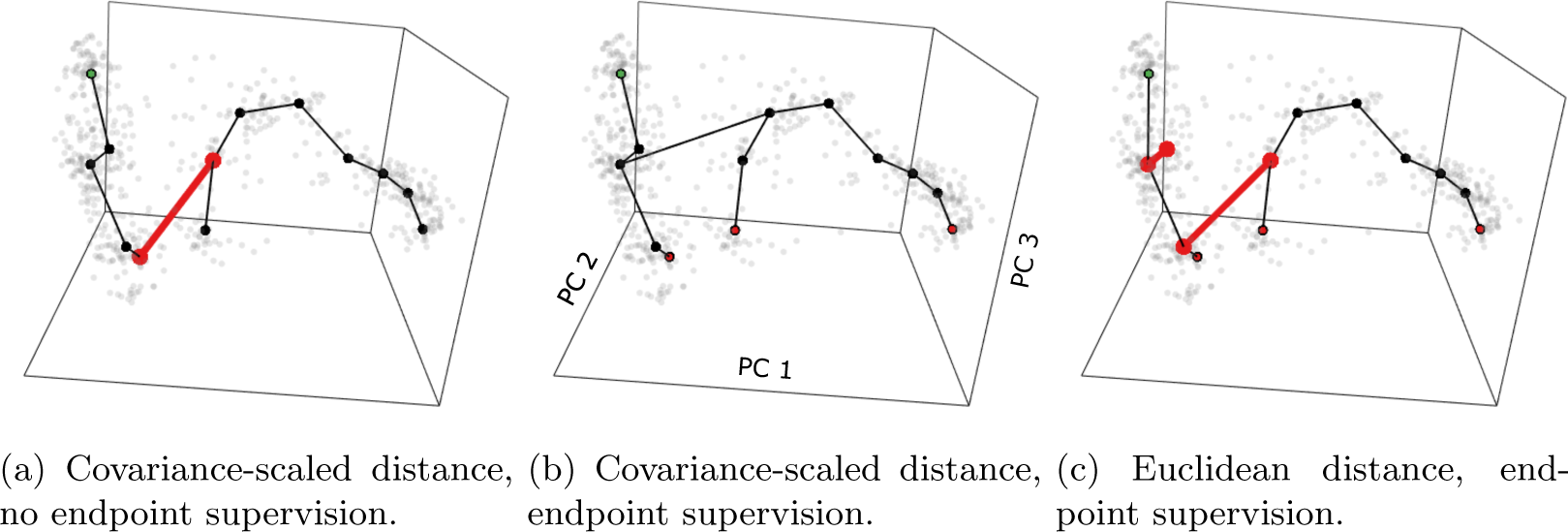
Semi-supervised lineage inference. We show three methods of constructing an MST on the clusters of the mouse olfactory epithelium dataset from Fletcher et al. (2017 in press). Although we only visualize the first 3 principal components, we note that these trees were constructed in the 5-dimensional space defined by the first 5 PCs. **Panel (a):** Without endpoint supervision, we draw the (known) false conclusion that sustentacular cells may develop into GBCs. **Panel (c):** Using Euclidean distances between cluster centers leads to the detection of a spurious branching event and the same erroneous conclusion about sustentacular cells. **Panel (b):** Both of these issues are resolved by using Slingshot’s covariance-scaled distance measure and marking the mature sustentacular cell cluster as an endpoint. This final tree is consistent with known biology and branching events were validated in follow-up experiments.

Other lineage detection methods were not able to correctly identify the three lineages in this dataset, diagrammed in Figure 5a. Monocle requires that the number of lineages be pre-specified, but in practice this number was unknown before running Slingshot and performing validation experiments. Even when we specified the proper number of lineages (three), Monocle’s results contradicted prior knowledge (Figure 5b). While one of Monocle’s lineages captured most of the neurogenic trajectory, it also included the identified microvillous cells and a fair number of immature sustentacular cells. More importantly, the neuronal lineage skipped most of the GBC cluster, cells which are known to be intermediates between HBCs and mature olfactory sensory neurons. Instead, Monocle classified GBCs as an alternate terminal state branching from the neuronal lineage. Another popular method, Wishbone, has a maximum of two lineages, so we applied it to a subset of the data believed to represent the two main lineages, namely, the sustentacular and neuronal lineages (Figures 5d and S4a). Wishbone identified a bifurcation, but not one which separated neurons from sustentacular cells (Figures 5e and S4b). The large skip in the longer lineage is caused by a small group of neurons which were isolated in a corner of the three-dimensional space constructed by diffusion maps. When we used Wishbone again with PCA as the dimensionality reduction step, we observed that this gap was no longer present, but Wishbone still failed to identify the primary bifurcation in the data (Figures 5f and S4b).

**Figure 5:**
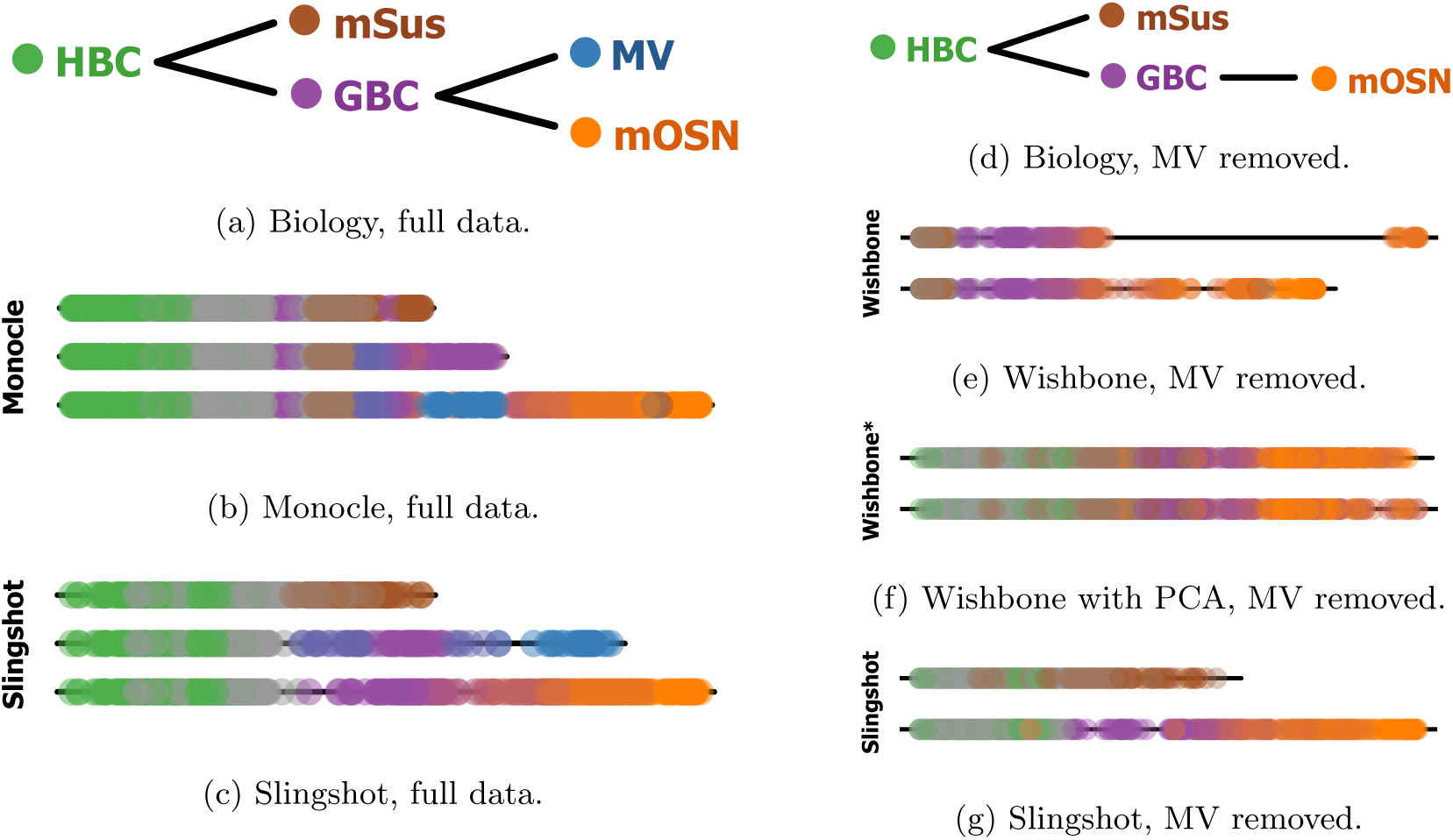
Multiple lineage inference. Results from multiple lineage inference methods on the olfactory epithelium dataset of Fletcher et al. (2017 in press), along with known biological relationships. **Panels (a–c):** Comparison of Slingshot and Monocle on the full dataset. For Slingshot, we specified the HBC cluster as the origin and the mature sustentacular (mSus) cluster as an endpoint. For Monocle, the main path was oriented such that pseudotimes started in the HBC cluster and the correct number of lineages (three) was provided. **Panels (d–g):** Comparison of Slingshot and Wishbone on a subset of the data, excluding cells specific to the microvillous (MV) lineage (we do this because Wishbone can only discern a maximum of two lineages).

For both the full and subsetted data, Slingshot identified lineages which are consistent with prior biological knowledge (Figures 5c and 5g). In the two-lineage case, this required no additional supervision beyond identifying the initial HBC cluster. For the full data, we specified all three terminal state clusters as endpoints, but we note that only one of these (the sustentacular cluster) had any effect on the inferred structure. Originally, it was not established that the microvillous endpoint was a branch off of the neuronal lineage, but this relationship was discovered and validated by Fletcher et al. (2017 in press). This novel biological discovery would not have been possible with lineage inference methods that restrict or require specification of the number of lineages.

In Supplemental Figures S3a and S3c, we show that in addition to capturing complicated, multilineage structures, Slingshot is also able to correctly detect a single lineage and two bifurcating lineages, respectively, in the datasets of Trapnell et al. (2014) and Shin et al. (2015). In both cases, Slingshot’s final pseudotime variables are highly similar to those found in the original papers, but do not rely on user specification of the number of lineages or subsetting of the data.

## 3 Discussion

We have introduced a new method, Slingshot, for lineage and pseudotime inference in datasets where the number of lineages is unknown. Because Slingshot breaks the lineage inference problem into two steps, we are able to make use of appropriate methods for each task and avoid the common trade-off between robustness and the flexibility to detect complex structures. Using a cluster-based MST for lineage identification allows Slingshot to identify potentially complex global patterns in the data without being overly sensitive to individual data points. Our novel simultaneous principal curves method for pseudotime inference extends the stability and robustness properties of principal curves to the case of multiple branching lineages.

Unlike other methods for multiple lineage identification, Slingshot does not require *a priori* knowledge of the number of lineages, but instead uses a cluster-based MST method that has the flexibility to detect novel lineages. At the same time, Slingshot also allows the user to constrain the global tree structure to ensure that previously established terminal states are correctly identified. This is a more natural way to incorporate prior information than placing burdensome restrictions on the number of lineages and leads to more biologically meaningful results than other methods. Lineage characteristics such as initial and terminal states can be difficult to identify at the level of individual cells (as in Wishbone), due to drop-out effects and the high levels of noise in single-cell data. However, more unsupervised methods like Monocle and Embeddr, which only require a direction to be given to an inferred lineage, can end up missing the initial state altogether. We find that supervision at the cluster level provides a nice balance: due to averaging, clusters are less ambiguous than individual cells, making them easier to identify based on known marker genes, and specifying initial and terminal states provides an intuitive, but not overly restrictive way to ensure that inferred lineages do not contradict previously established results.

We demonstrated, using the data of Trapnell et al. (2014) and Shin et al. (2015), that Sling-shot can correctly detect a single lineage and two bifurcating lineages, respectively (Supplemental Figures S3a and S3c). In both cases it produces results similar to those found and validated in the original paper (Supplemental Figures S3b and S3d). Additionally, using the olfactory epithelium data of Fletcher et al. (2017 in press), we demonstrated that with minimal supervision, Slingshot can correctly identify a complicated three-lineage structure that other methods cannot.

Since there are many aspects to the problem of lineage inference, from sample collection to final analysis, it is important to define precisely the tasks for which Slingshot is designed. The philosophy of Slingshot is that some common steps in single-cell analysis, such as dimensionality reduction or clustering, do not have a single solution that works well for all data types. For example, Slingshot does not require a specific dimensionality reduction method because single-cell data can come from a variety of assays and in a wide range of dimensions, from the 271 cells × 47, 192 genes RNA-Seq dataset of Trapnell et al. (2014) to the 25, 000 cells × 13 markers mass cytometry dataset of Setty et al. (2016). This extreme heterogeneity precludes any one-size-fits-all solution and similar arguments can be made for other upstream analysis steps. Slingshot makes use of both a dimensionality reduction and clustering step, but unlike other methods it does not attempt to solve either problem, viewing these instead as separate analysis choices, analogous to the choice of normalization method or short read aligner.

Ultimately, single-cell data are noisy, high-dimensional, and may contain a multitude of competing, interwoven signals. In the presence of such data, Slingshot provides a robust and modular method for lineage inference that allows for novel lineage discovery, meaningful incorporation of biological constraints, and fits easily within existing analysis pipelines.

## 4 Online Methods

We start from an *n × J* matrix of normalized expression measures (e.g., read counts) for *n* single cells and *J* genes or features. Slingshot assumes that the *n* cells have been partitioned into *K* clusters, potentially corresponding to distinct cellular states. Although Slingshot can in principle be applied directly to the normalized expression values, we strongly recommend a dimensionality reduction step before pseudotemporal reconstruction, as Slingshot’s curve-fitting step uses Euclidean distances which can misbehave in high-dimensional spaces (cf. curse of dimensionality). Dimensionality reduction can also strengthen signal in the data and help with visualization. We denote the dimension of the reduced space by *J* ^*′*^.

Before detailing Slingshot’s two main steps, we introduce some notation. First, denote by *X* = (*X*_*ij*_) the *n × J ′* reduced-dimensional matrix of gene expression measures, for cells *i ϵ* {1, …, *n*} and dimensions *j ϵ* {1, …, *J′*}. Let {*C*_1_, …, *C*_*K*_} denote the *K* cell clusters or states, i.e., disjoint subsets of cells, obtained by clustering the cells based on their gene expression measures. We then define a lineage as an ordered set of clusters and let *L* denote the total number of lineages. For a particular lineage, *L*_*l*_, denote its length (i.e., the number of clusters in the lineage) by *K*_*l*_ and the *k*^th^ cluster by 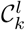, for *l ϵ {*1, …, *L*} and *k ϵ {*1, …, *K*_*l*_}. In particular, 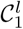 and 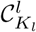 correspond to the initial and terminal states for the *l*^th^ lineage, respectively. It is important to note that a cluster can belong to multiple lineages and that the ordering of the clusters within a lineage does not strictly determine the final relative orderings of cells in those clusters.

As a given cluster can belong to multiple lineages, so can a cell. For ease of notation, we therefore allow cells to have distinct pseudotime values for each lineage they are a part of. The pseudotime value for cell *i* in lineage *l* is denoted by 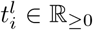; if cell *i* does not belong to lineage *l*, i.e., 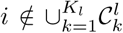, then set 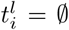. The vector of pseudotime values for lineage *l* is denoted by 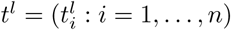.

### 4.1 Identification of Cluster-Based Lineages

In its first step, Slingshot identifies lineages by treating clusters of cells as nodes in a graph and drawing an MST between the nodes, similar to the work of Ji and Ji (2016) and Shin et al. (2015). Lineages are then defined as ordered sets of clusters created by tracing paths through the MST, starting from a given root node. Our method differs however in a number of important respects from those of Ji and Ji (2016) and Shin et al. (2015), including the distance measure used for drawing the tree and the incorporation of biologically meaningful supervision.

#### 4.1.1 Shape-Sensitive Distance Measure between Cell Clusters

Constructing an MST involves specifying a distance measure between nodes (in this case, clusters). Although in principle any type of distance measure could be used (e.g., Euclidean, Manhattan), we found that a Mahalanobis distance, i.e., a covariance-scaled Euclidean distance, that accounts for cluster shape, works well in practice. Specifically, the pairwise distance between clusters *i* and *j*, *d*(*C*_*i*_*, C*_*j*_), is defined as

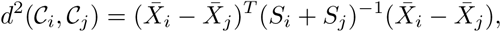

where 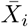 represents the center (mean) of cluster *i* and *S*_*i*_ its empirical covariance matrix in the reduced-dimensional space. This is essentially a multivariate *t*-statistic. By default, Slingshot uses the full covariance matrix of both clusters, allowing us to draw trees that are better covered by and representative of the cells in a dataset. However, in the presence of small clusters, the matrix *S*_*i*_ + *S*_*j*_ may not be invertible and we replace the full covariance matrix with the corresponding diagonal covariance matrix.

#### 4.1.2 Biologically Meaningful Supervision

Slingshot allows two forms of supervision during lineage identification: initial state and terminal states specification. Like other methods relying on cluster-based MSTs (TSCAN, Waterfall), Slingshot requires the user to identify the initial cluster or root node. Subsequently, every direct path from this node to a leaf node (i.e., a cluster with only one edge) will be called a lineage. Indeed, all existing lineage construction methods explicitly or implicitly make the assumption that a starting state can be identified by the user: Monocle and Embeddr construct orderings for which the user must select the correct direction and Wishbone requires the user to select an initial cell or group of cells. In the simple case where the MST constructed by Slingshot has only two leaf nodes and one is specified as the root, this results in a single lineage. If an interior (non-leaf) node is specified as the origin, this results in two lineages, one terminating in each leaf node. Clusters with more than two edges will create bifurcations and produce additional lineages.

Additionally, Slingshot allows the user to provide further supervision in the construction of the lineages by selecting clusters known to represent terminal cell states, imposing a constraint on the MST algorithm. The constrained MST is obtained by first constructing the MST on all *non-selected* clusters and then connecting each *selected* cluster to its nearest non-selected neighbor. Such supervision results in more biologically meaningful lineages for situations where the data can be explained by many possible lineage structures. Identified lineages are by construction consistent with known biology and provide improved stability over less supervised methods. Although terminal state supervision is not required, we find that in many settings researchers do have knowledge of the cell types present in their data and that systematically incorporating this knowledge can provide more stable inference. Ultimately, detecting multiple lineages based on gene expression is a difficult problem that benefits from such guidance, as we demonstrate in the results (Section 2).

### 4.2 Identification of Individual Cell Pseudotimes

The second stage of Slingshot is concerned with assigning pseudotimes to individual cells. For this purpose, we make use of principal curves (Hastie and Stuetzle, 1989) to draw a path through the gene expression space of each lineage. As we show in the results (Section 2), principal curves give very robust pseudotimes when there is a single lineage.

Multiple lineages demand more care and are handled using the simultaneous principal curves method proposed below. Just as clusters in the MST may belong to one or more lineages, the cells which constitute these clusters may be assigned to one or more lineages. In principle, we could construct traditional principal curves for each lineage separately to arrive at pseudotimes. However, there is no guarantee that these curves would agree with each other in the neighborhood of clusters shared between lineages, so cells belonging to multiple lineages could be assigned very different pseudotime orderings by each curve. Since we assume a smooth differentiation process, this is potentially a violation and may be problematic in downstream analysis.

We therefore introduce a method of simultaneously fitting the principal curves of each lineage, which shrinks the curves to a consensus path in areas where lineages share many common cells, but allows the curves to separate as they share fewer and fewer cells. This ensures smooth bifurcations of the paths. We call the resulting curves *simultaneous principal curves*, as they are fit by an iterative procedure based on the principal curves algorithm of Hastie and Stuetzle (1989). When there is only a single lineage (*L* = 1), the pseudotimes of Slingshot are found by the standard principal curves algorithm, except that the initial curve for the algorithm is based on the lineage’s path through the MST found in the earlier stage (see below for details), rather than the first principal component.

We review the standard principal curves algorithm (for a single curve) in order to be clear about how we adapt it for simultaneous principal curves. After specification of an initial curve, the algorithm iteratively follows these steps:

1. Project all data points onto the curve and calculate the arclength from the beginning of the curve to each point’s projection. Setting the lowest value to zero, this produces pseudotimes.
2. For each of the *J′* dimensions, use the cells’ pseudotimes to predict their coordinates along dimension *j*, *j ϵ* {1, …, *J′*}, typically with a smoothing spline. This produces a set of functions which collectively map pseudotime values in ℝ_≥0_ into ℝ*J′*, thereby defining a smooth curve in *J′* dimensions.
3. Repeat this process until convergence. We use the sum of squared distances between cells and their projections on the curves to determine convergence.

In the case of branching lineages, this iterative process is modified with an additional shrinkage step to ensures smooth bifurcations. Specifically, for each lineage *l*, we infer a vector of pseudotime values, 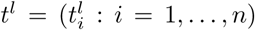, and a function *c*_*l*_: ℝ_≥0_ ⟶ ℝ^*J′*^ for the associated curve in the low-dimensional space. Then, the shrinkage step is done by first constructing an average curve which, as with the individual lineage curves, is a function of pseudotime. It simply consists of the average of the points along the *L* curves at each pseudotime value:

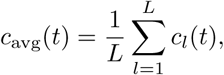

where *t* represents pseudotime. Because the *L* lineages share the same root cluster, we are assured that their starting points (where *t* = 0) will be identical and will also be the starting point of the average curve.

We next construct a set of lineage-specific weighting functions to determine how much we should shrink each curve toward the average. For lineage *l*, we define the weighting function *w*_*l*_: ℝ_≥0_ *⟶* [0, 1], with the constraint that *w*_*l*_ must be non-increasing. Additionally, by specifying that *w*_*l*_(0) = 1*, ∀l*, we ensure that diverging curves always share the same initial point. These weighting functions allow us to shrink the diverging lineage curves toward their shared average curve with the additional update step:

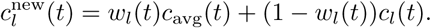

If all the *w* functions are smooth, this shrinkage step ensures that the final curves will follow a tree structure with smooth branching events.

Slingshot’s default weighting function satisfies these conditions and is based on the distribution of pseudotimes over cells shared between lineages. Given a set of lineages *l*_1_, …, *l*_*m*_ (typically with *m* = 2), which all contain certain shared cells 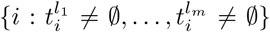, we define the weighting function for lineage *l*_1_ as follows. Set 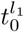 and 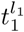 respectively as the lowest and highest non-outliers in this distribution, where outliers are defined by the 1.5IQR rule of boxplots. The weighting function is then defined as:

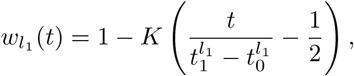

where *K* is the cumulative distribution function of a standard cosine kernel with a bandwidth of 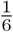 (which places weight only on values between 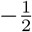 and 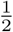). We note that 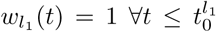 and 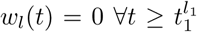. Weighting functions for the other lineages are then calculated similarly. The final curves are fairly robust to the choice of kernel function (we also tried standard kernels provided by the density function in R, results not shown).

In both the single and branching lineage cases, final pseudotime values are derived from each point’s orthogonal projection onto the curves. Thus, cells belonging to multiple lineages will have multiple pseudotime values, but these values will agree quite well for cells positioned before the lineage bifurcation, where the curves are most similar.

#### Initialization of Simultaneous Principal Curves Algorithm

As mentioned above, we initialize the algorithm using the MST from the first stage. Specifically, we start with the piecewise linear path through the centers of the clusters contained in the lineage (in contrast, standard principal curves are initialized by the first principal component of all points being fit). Starting with the path through the cluster centers allows us to leverage the prior knowledge that went into lineage identification as well as to improve the speed and stability of the algorithm, though in practice, the two procedures typically converge to very similar curves.

### 4.3 Datasets

We demonstrate the performance of Slingshot by applying it to three previously-published single-cell RNA-Seq datasets, each with a different number of terminal cell types. The first is a subset of the data used in Trapnell et al. (2014), which assayed 271 human skeletal muscle myoblasts (HSMMs) in order to study their development into mature myotubes. This is an example of data with only a single lineage. In their analysis, Trapnell et al. (2014) identify a population of contaminating interstitial mesenchymal cells, which we omit from this dataset. This results in a sample of 212 cells believed to form a single, continuous developmental lineage. For our analysis, we used the cluster labels generated by Monocle as well as a set of labels obtained via *k*-means clustering and, as in the original paper, we represented the data in two dimensions obtained by ICA. The normalized data were downloaded from the NCBI GEO database (accession GSE52529).

The second dataset comes from Shin et al. (2015), who assayed 132 hippocampal quiescent neural stem cells (qNSCs) and their immediate progeny from adult mice, cells known to be involved in neurogenesis. Their goal was to assess cellular heterogeneity among this population and analyze continuous-time developmental dynamics. After removing a few outliers, their analysis focuses on 101 cells believed to represent the development of qNSCs into intermediate progenitor cells (IPCs), a transitional state between qNSCs and mature neurons. However, they note an additional cluster of 23 cells branching off of this lineage, potentially representing an alternative terminal cell type. As in the original paper, we used the hierarchical clustering labels and the first two principal components as the reduced dimensional space. Rather than focus solely on the IPC lineage though, we characterized the developmental trajectory of both proposed cell fates. The normalized data and code for preliminary analysis were downloaded from GEO (accession GSE71485).

The third dataset is that of Fletcher et al. (2017 in press), featuring 616 cells from the adult mouse olfactory epithelium (OE), tracing the development of quiescent stem cells into three unique terminal cell fates. The primary lineage maps the development of horizontal basal cells (HBCs) into mature olfactory sensory neurons (OSNs) via a series of intermediate states. The secondary lineage gives rise to the support (sustentacular) cells of this system and features fewer identifiable intermediates. A third lineage which appears to split from the neuronal path leads to a cluster of microvillous cells. Again, we relied on the cluster labels of the authors, who used RSEC (Purdom and Risso, 2016), and represent cells by their coordinates along the first five principal components. The normalized data and cluster labels are available from GEO (accession GSE95601).

## 5 Declarations

### 5.1 Ethics Approval and Consent to Participate

Ethical approval and consent were not applicable to this study.

### 5.2 Availability of Data and Material

Slingshot is implemented in the R programming language and publicly available through an open-source package on GitHub (https://github.com/kstreet13/slingshot).

### 5.5 Authors’ Contributions

K.S., D.R., E.P., and S.D. formulated the methodology. J.N. participated in experimental strategy and design. N.Y. conceived the endpoint supervision strategy. K.S. and D.R. wrote the R package. K.S., R.B.F., and D.D. performed the data analysis. K.S., D.R., E.P., and S.D. wrote the manuscript.

## 5.6 Acknowledgements

We thank Mary Combs for suggesting the name Slingshot.

